# NG2 glia are required for maintaining microglia homeostatic state

**DOI:** 10.1101/667402

**Authors:** Yingjun Liu, Adriano Aguzzi

## Abstract

Microglia play vital roles in the health and diseases of the central nervous system. Loss of microglia homeostatic state is a key feature of brain aging and neurodegeneration. However, the mechanisms underlying the maintenance of distinct microglia states are largely unclear. Here we show that NG2 glia, also known as oligodendrocyte precursor cells, are essential for maintaining the homeostatic microglia state. We developed a highly efficient and selective NG2 glia depletion method using small-molecule inhibitors of platelet-derived growth factor signaling in cultured brain slices. We found that loss of NG2 glia abolished the homeostatic microglia signature without affecting the disease-associated microglia profiles. Similar findings were also observed *in vivo* by genetically ablating NG2 glia in the adult mouse brain. These data suggest that NG2 glia exert a crucial influence onto microglia cellular states that are relevant to brain aging and neurodegenerative diseases. In addition, our results provide a powerful, convenient and selective tool for the investigation of NG2 glia function.

**Main points:**

1. Postnatal NG2 glia maintenance obligatorily depends on continuous PDGF signaling.
2. A highly efficient, selective and versatile NG2 glia depletion method is established.
3. Loss of NG2 glia abolishes the homeostatic microglia signature both *in vitro* and *in vivo*.

## Introduction

Microglia are innate immune cells residing in the brain, playing critical roles in the development, maturation, maintenance, aging and functional deteriorations of the central nervous system (CNS). The cellular activities of microglia are diverse, ranging from protective to detrimental depending on cellular “states” defined by the characteristic expression of different sets of signature genes (Keren-Shaul et al. 2017; Krasemann et al. 2017). In the healthy CNS, microglia exhibit a distinct resting or homeostatic state featured by the expression of genes that are vital for the defensive activities of microglia against various disease-associated events (Kierdorf and Prinz 2017). Upon brain injuries and in the aging brain, or in brains affected by neurodegenerative diseases, the homeostatic state of microglia is lost, accompanied by the induction of disease-associated phenotypes (Butovsky and Weiner 2018; Keren-Shaul et al. 2017; Krasemann et al. 2017; Zrzavy et al. 2017). The transition of microglia state from homeostatic to disease-associated is crucial for the development of neurodegenerative disorders and reversing such transition holds great potential for the restoration of normal microglia function and intervention of the disease.

Accumulating evidence suggest that the regulation of microglia phenotypes is a highly localized event and microglia with distinct cellular states co-exist in the aged brains or in brains affected by neurodegenerative diseases (Liu and Aguzzi 2019; Sala Frigerio et al. 2019). In Alzheimer’s disease (AD), for example, microglia associated with amyloid beta (Aβ) plaques show drastic downregulation of the homeostatic signature genes compared with microglia in adjacent brain regions (Keren-Shaul et al. 2017; Krasemann et al. 2017), while a subset of microglia in the AD brain show a strong pro-inflammatory phenotype characterized by inflammasome activation (Heneka et al. 2013). The mechanisms underlying the localized regulation of microglia cellular phenotypes are still unclear.

NG2 glia, also known as oligodendrocyte precursor cells, are a distinct glial cell population distributed throughout the adult CNS. They represent 5–10% of the total cell population in the brain. NG2 glia are identified by the co-expression of NG2 (also known as chondroitin sulfate proteoglycan 4) and platelet-derived growth factor receptor alpha (PDGFRα), and function as oligodendrocyte precursor cells during development or myelin damages (Kang et al. 2010; Tripathi et al. 2010; Trotter et al. 2010). Interestingly, in contrast to microglia, a recent study found that Aβ plaque-associated NG2 glia in the AD brain showed a senescent phenotype (Zhang et al. 2019), suggesting the impaired NG2 glia function in the plaque-containing microenvironment. Whether the local loss of NG2 glia function contributes to the disturbed homeostatic microglia signature around Aβ plaques or more generally, whether NG2 glia play a role in maintaining microglia homeostatic state in the healthy CNS is unknown.

Here, we developed a selective and efficient NG2 glia depletion method through inhibiting the platelet-derived growth factor (PDGF) signaling with small chemical molecules. We found that loss of NG2 glia abolished the microglia homeostatic signature in the cultured brain slices and *in vivo*, revealing a critical role of NG2 glia in the regulation of microglia homeostatic state. This newly identified NG2 glia-microglia interaction may have broad implications for the physiology and pathophysiology of the brain.

## Results

### PDGFR inhibitor induces highly efficient NG2 glia depletion

Absence of PDGF signaling impairs NG2 glia generation during embryonic development (Calver et al. 1998; Fruttiger et al. 1999) and disturbs NG2 glia maintenance in primary cultures (Barres et al. 1992; Barres et al. 1993). Therefore, we hypothesized that manipulating PDGF signaling might modify NG2 glia abundance and function. To test this, we treated cultured brain slices derived from different brain regions with two potent PDGFR inhibitors: CP673451 and Crenolanib. Western blotting examination showed a strong decrease of the protein levels of PDGFRα and NG2, the markers for NG2 glia, in both CP673451 and Crenolanib treated slices (**Fig. 1a-b**). A similar treatment did not reduce the expression of either gene in the oligodendrocyte precursor cell line Oli-neu (**Supplementary fig. 1a-b**), suggesting that the decreased PDGFRα and NG2 protein levels in the CP673451 and Crenolanib treated brain slices were resulted from the loss of NG2 glia rather than from inhibiting PDGFRα and NG2 gene expression. Indeed, through immunofluorescence, we observed a drastic reduction of both PDGFRα^+^ and NG2^+^ cell numbers in the CP673451 and Crenolanib treated slices (**Fig. 1c-d**; **Supplementary fig. 1c-g**). Time- and dose-dependency analyses indicated that 7 days of treatment at 500 nM concentration could achieve >95% NG2 glia depletion for both inhibitors (**Fig. 1e-h**). These results suggest that PDGF signaling inhibitors may represent universal tools for efficient NG2 glia depletion.

**Fig 1.**
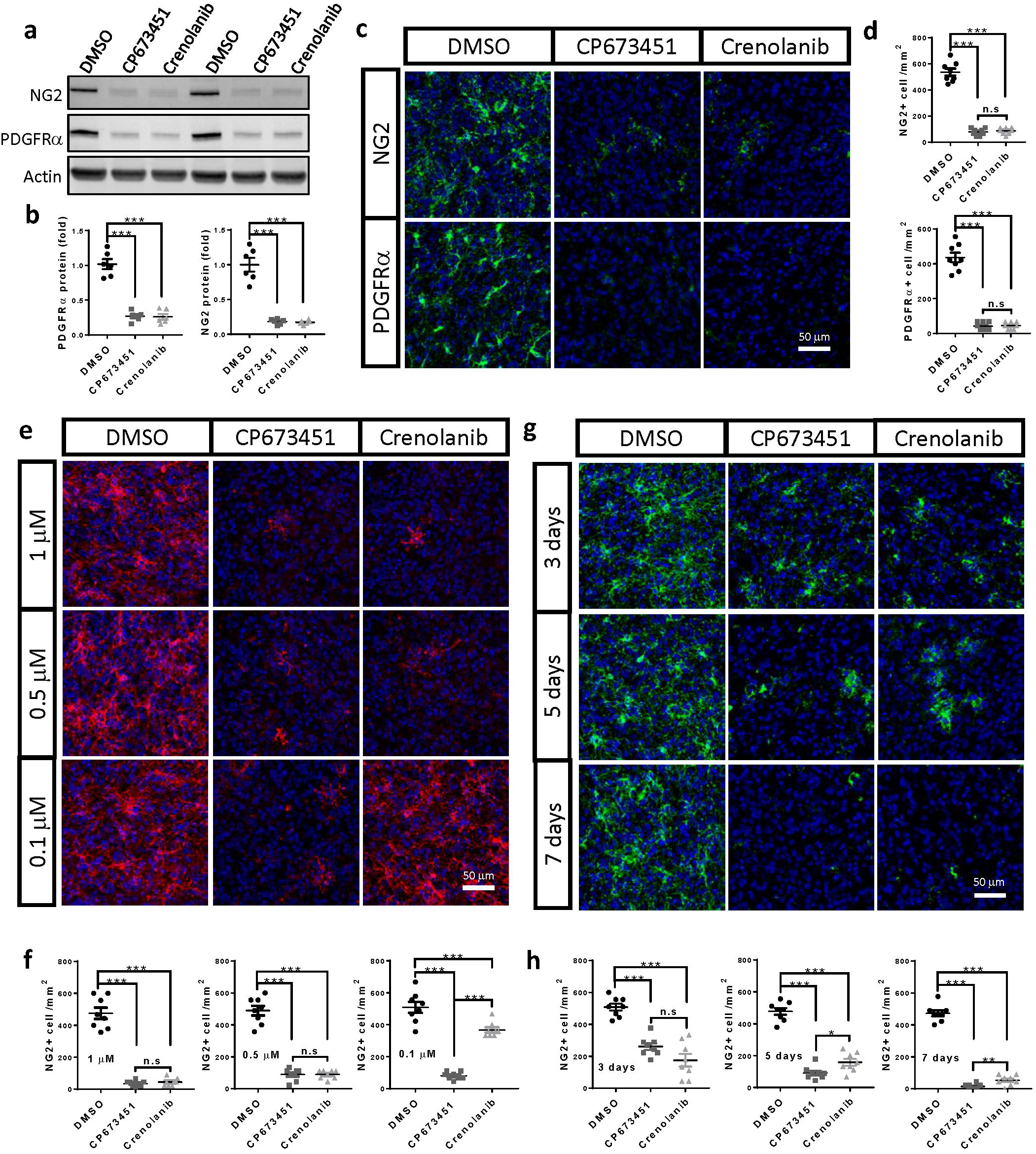
PDGFR inhibitors induce highly efficient NG2 glia depletion. **a-b**, western blots (a) and quantification (b) of PDGFRα and NG2 in slices treated with CP673451 and Crenolanib for 9 days. **c**, immunofluorescence of PDGFRα and NG2 on slices treated with CP673451 and Crenolanib for 9 days. Scale bar: 50 μm. **d**, quantification of PDGFRα^+^ and NG2^+^ cell numbers shown in **c. e**, immunofluorescence of NG2 on slices incubated with 1 μM, 500 nM and 100 nM CP673451 or Crenolanib for 9 days. Scale bar: 50 μm. **f**, quantifications of NG2^+^ cell numbers shown in **e. g**, immunofluorescence of NG2 on slices incubated with 500 nM CP673451 and Crenolanib for 3, 5 and 7 days. Scale bar: 50 μm. **h**, quantifications of NG2^+^ cell numbers shown in **g**. * P<0.05; ** P<0.01; *** P< 0.001. n.s, not significant.

### PDGFR inhibitor induced NG2 glia depletion is highly specific

To determine the specificity of our NG2 glia depletion approach, we examined the numbers of various cell types present in the adult CNS by immunofluorescence after CP673451 and Crenolanib treatment. We found that PDGFR inhibitors did not affect the numbers of neurons, astrocytes, oligodendrocytes and microglia, nor did they affect the amount of myelin or elicit microglia activation and inflammatory responses (**Fig. 2a-b**; **Supplementary fig. 2a-c**). Previous studies have suggested that pericytes express the beta form of PDGFR (PDGFRβ). We wondered whether the treatment affected the number of pericytes. Interestingly, we found that an anti-PDGFRβ antibody from Cell Signaling Technologies recognized two distinct populations of cells (**Fig. 2a**): one with the classical morphology of pericytes, and the other with the morphology of NG2 glia (possibly due to cross reactivity with PDGFRα). However, almost all the PDGFRβ^+^ cells with NG2 glia morphology disappeared after the inhibitor treatment, while the number of PDGFRβ^+^ cells with pericyte morphology was unchanged (**Fig. 2a-b**). Similarly, we found there was no detectable difference between groups regarding the number of CD13^+^ pericytes (**Fig. 2a-b**). These results suggest that PDGFR inhibitors induce NG2 glia depletion with high cell-type selectivity.

**Fig 2.**
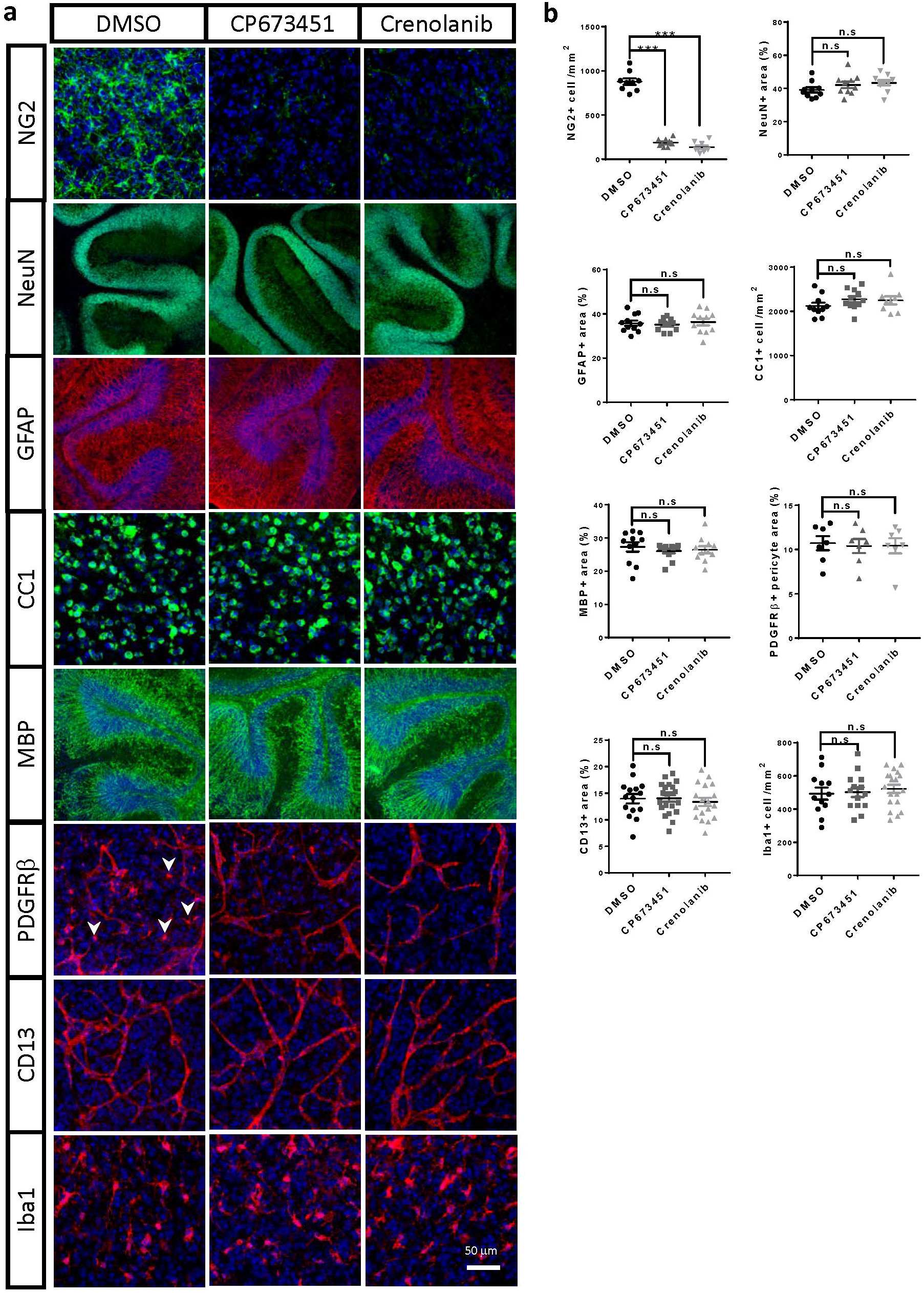
PDGFR inhibitor-based NG2 glia depletion is highly cell-type specific. **a**, immunofluorescence of NG2, NeuN, GFAP, CC1, MBP, PDGFRβ, CD13 and Iba1 on slices treated with 500 nM CP673451 and Crenolanib for 9 days. Scale bar: 50 μm. Arrowheads: PDGFRβ^+^ cells with NG2 glia morphology. **b**, quantifications of cell numbers or signal positive areas shown in **a**. *** P< 0.001. n.s, not significant.

### Loss of NG2 glia abolishes the homeostatic microglia signature

To identify any functional consequences of NG2 glia depletion, we performed transcriptome analyses of cultured brain slices treated with CP673451, Crenolanib, or DMSO as control. By setting a cut-off of *p* < 0.01 and log2 ratio > 0.5, we found 459 and 464 differentially expressed genes (DEGs) in CP673451 and Crenolanib-treated slices compared with controls, respectively (**Fig. 3a; Supplementary table 1; Supplementary table 2**). In contrast, only 19 genes were differentially expressed between CP673451 and Crenolanib-treated slices (**Supplementary fig. 3a**), suggesting that the two inhibitors produced very similar biological effects. We isolated the genes that differentially expressed in both CP673451 and Crenolanib treated groups compared with controls and assigned these genes as core-DEGs (**Fig. 3b**; **Supplementary fig. 3b**; **Supplementary table 3**).

**Fig 3.**
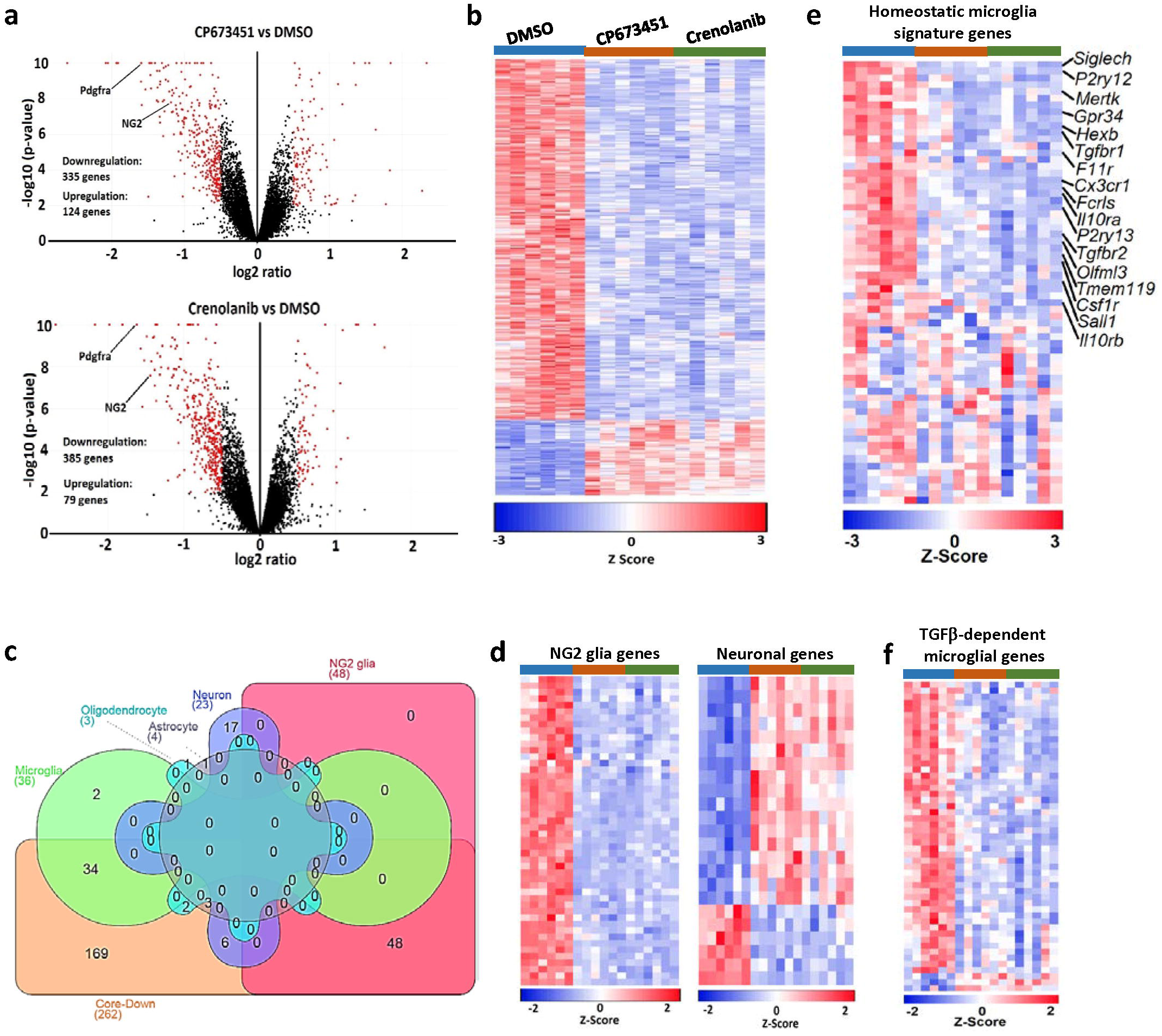
Absence of NG2 glia disturbs microglia homeostatic state. **a**, volcano plots of RNAseq data comparing CP673451-treated (upper image) or Crenolanib-treated (lower image) with DMSO-treated slices. **b**, heatmap of genes differentially expressed between both CP673451 vs. DMSO and Crenolanib vs. DMSO (core-DEGs). **c**, assignment of core-DEGs to specific cell types according to published cell-type enrichment datasets. Core-Down: downregulated genes within the core-DEG collection. **d**, heatmaps of overlapping genes between the core-DEGs and NG2 glia-enriched (left image) or neuron-enriched genes (right image). **e**, heatmap of the 65 homeostatic microglia signature genes after NG2 glia depletion. **f**, heatmap of significantly dysregulated TGFβ-dependent microglia genes after NG2 glia depletion.

To determine which cell types were affected by NG2 glia depletion, we examined the overlaps between the core-DEGs and the cell-type enriched genes extracted from a previous study (Zhang et al. 2014). We found that 48 out of the 436 NG2 glia-enriched genes were present in the core-DEGs (**Fig. 3c**). Remarkably, all of these 48 genes were downregulated after CP673451 and Crenolanib treatment (**Fig. 3d**), further confirming the efficient depletion of NG2 glia after PDGFR inhibitor treatment. In contrast, there were only three and four overlapping genes between the core-DGEs and the 385 oligodendrocyte and 704 astrocyte enriched genes, respectively (**Fig. 3c**). Twenty three neuron-enriched genes were present in the core-DEGs, of which 17 were upregulated and six were downregulated (**Fig. 3c-d**). Interestingly, we identified 36 overlaps between the core-DEGs and microglia-enriched genes, 34 of which were downregulated (**Fig. 3c**). Some of the downregulated microglia genes, such as Tmem119 and Olfml3, have been reported to be highly expressed in homeostatic microglia (Bennett et al. 2016; Butovsky et al. 2014). Therefore, we extracted the expression profiles of 65 previously identified homeostatic microglia signature genes (Krasemann et al. 2017) from our dataset and found that around half of these genes were downregulated in both CP673451-treated and Crenolanib-treated brain slices (**Fig. 3e**). The downregulation of these genes cannot be attributed to direct effects of CP673451 and Crenolanib on microglia because microglia do not express PDGFRs and, more importantly, CP673451 and Crenolanib treatment had no influence on the expression of these genes in cultured BV2 microglia (**Supplementary fig. 3c**). Since microglia numbers were unchanged after NG2 glia depletion (**Fig. 2a-b**), these results suggest that NG2 glia are required for the maintenance of microglia homeostatic state. In line with the findings that loss of NG2 glia did not induce microglia activation and inflammatory responses (**Supplementary fig. 2**), we found there were no overall expressional changes of genes that represent disease-associated microglia states after NG2 glia depletion (**Supplementary fig. 3d**).

### Absence of NG2 glia impairs the microglia TGFβ signaling

Previous studies indicated that TGFβ signaling played vital roles in maintaining microglia homeostasis (Butovsky et al. 2014; Qin et al. 2018; Zoller et al. 2018). Interestingly, we found both the type-1 and type-2 TGFβ receptors, which are highly expressed in homeostatic microglia, were notably downregulated after NG2 glia depletion (**Fig. 3e**). In addition, the expression of latent TGFβ binding protein 2 (LTBP2), an important regulator for TGFβ secretion and activation, was dramatically decreased by CP673451 and Crenolanib treatments (**Supplementary fig. 3e**). These results suggest that the TGFβ signaling might be impaired in the NG2 glia-depleted slices. We therefore examined the expression profiles of 112 TGFβ-dependent microglia genes identified previously (Butovsky et al. 2014) in our RNAseq dataset. Indeed, we found that 48 of them downregulated after removing NG2 glia (**Fig. 3f**). These data supports a notion that NG2 glia depletion causes microglia to lose their homeostatic state by suppressing the TGFβ signaling.

### NG2 glia are required for maintaining microglia homeostatic signature in vivo

Next, we set out to determine whether NG2 glia are required for the maintenance of microglia homeostatic state *in vivo*. We generated a mouse line that expresses diphtheria toxin receptor (DTR) selectively in NG2 glia by crossing the PDGFRα-CreER™ mice (Kang et al. 2010) with the inducible DTR (iDTR) mice (Buch et al. 2005). We induced DTR expression in NG2 glia by feeding two-month old mice with tamoxifen-containing diet (**Fig. 4a**). One month after terminating the tamoxifen treatment, we administered diphtheria toxin (DT) for five consecutive days and collected the brains 3 days later (**Fig. 4a**). Western blotting and immunofluorescence confirmed a reduction of NG2 glia numbers by approximately 50% after DT injection in multiple brain regions (**Fig. 4b-c**; **Supplementary fig. 4a-e**). No significant changes of cell-type markers were found for neuron, astrocyte, microglia, oligodendrocyte and pericyte after DT injection (**Supplementary fig. 4f-g**). Similar to the findings from cultured slices, suppression of NG2 glia conspicuously reduced the expression of homeostatic-microglia genes such as Tmem119 and Olfml3 in adult mice (**Fig. 4d**). In contrast, there was no significant change of CD11b (**Fig. 4d**), a maker gene for microglia at different cellular states. These results suggest that loss of NG2 glia disturbs microglia homeostatic state without influencing microglia abundance *in vivo*. Immunofluorescent examinations further supported this conclusion. We found there were no changes of Iba1^+^ microglia number and Iba1 immuno-intensity in the brain after NG2 glia depletion (**Fig. 4e-f**). In contrast, the immuno-intensity of Tmem119 but not the Tmem119^+^ microglia number was profoundly decreased (**Fig. 4e-f**), suggesting a downregulation of Tmem119 in microglia in the NG2 glia depleted brain, which is consistent with the qRT-PCR results (**Fig. 4d**). As in cultured brain slices, we found no difference in the expression of inflammatory markers between the NG2 glia-depleted and control brains (**Supplementary fig. 4h**). All of these results confirm that the absence of NG2 glia disturbs the homeostatic state of microglia without inducing disease-associated microglia phenotypes or an inflammatory response in the adult mouse brain.

**Fig 4.**
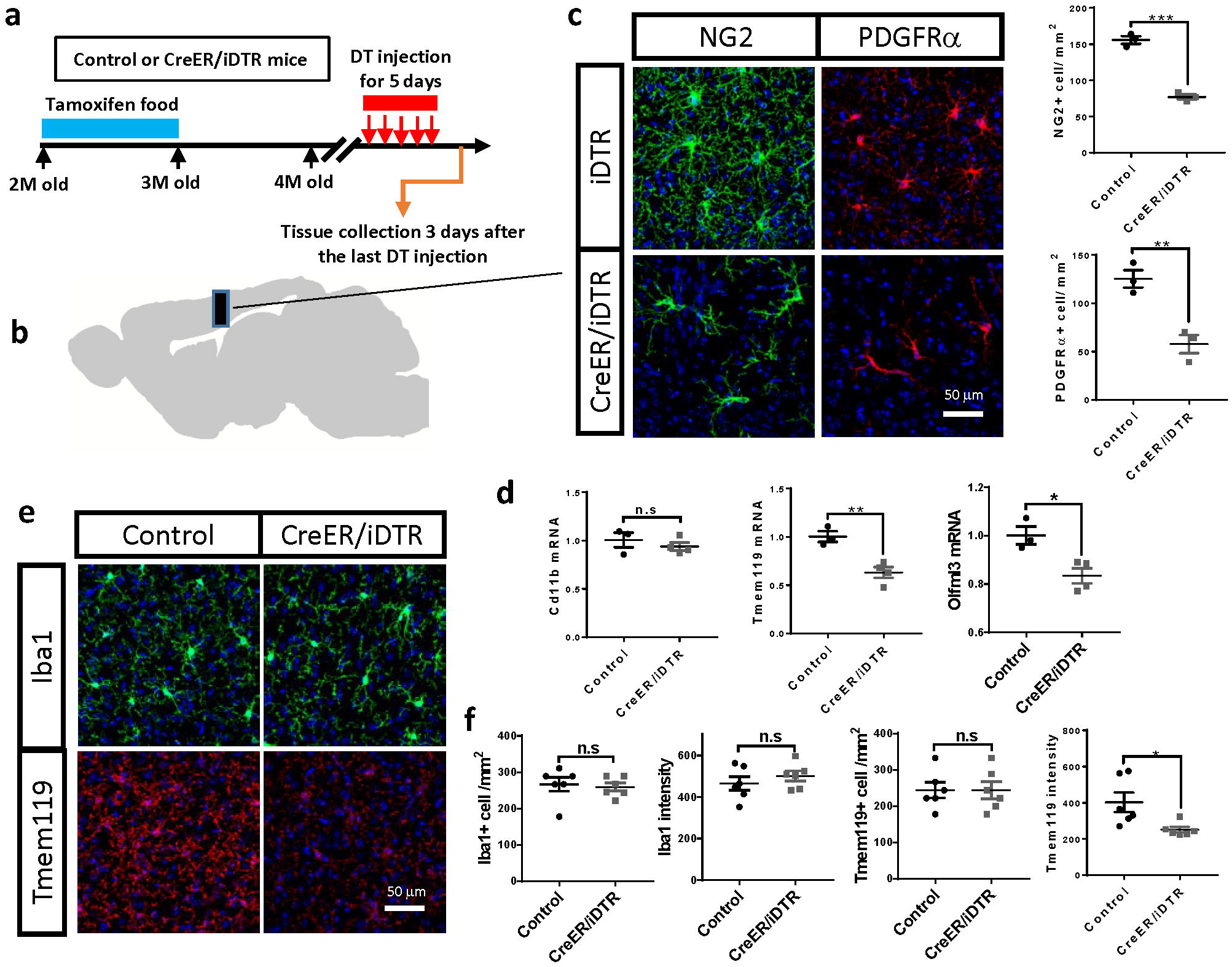
Loss of NG2 glia downregulates homeostatic microglia signature *in vivo*. **a**, experimental strategy of NG2 glia depletion in the adult mice. **b-c**, representative immunofluorescent images and quantification of PDGFRα^+^ and NG2^+^ cells in the brains of NG2 glia-depleted and control mice. Scale bar: 50 μm. **d**, qRT-PCR results of microglia homoeostatic genes after NG2 glia depletion in the adult mouse brain. **e**, representative images of Iba1 and TMEM119 immunostaining in the NG2 glia-depleted and control brains. Scale bar: 50 μm. **f**, quantifications of Iba1^+^ and Tmem119^+^ microglial cell number as well as Iba1 and Tmem119 immuno-intensity shown in **e**. * P<0.05; ** P<0.01; *** P< 0.001. n.s, not significant.

## Discussion

Although NG2 glia were first described more than three decades ago (Stallcup 1981), their role in brain homeostasis and disease remains elusive, partly because there are few tools for the selective and efficient manipulation of NG2 glia. Here, we show that tyrosine-kinase inhibitors interfering with the PDGF pathway can deplete NG2 glia efficiently and selectively. Using these tools, we discovered that NG2 glia control the transition of microglia into specific transcriptional states. While NG2 glia depletion induced a loss of the homeostatic profile of microglia, it did not induce a disease-associated microglia phenotype (Keren-Shaul et al. 2017; Krasemann et al. 2017) or a neuroinflammatory response. Therefore, the transcriptional profile of microglia in the absence of NG2 glia appears to represent a phenotypic state that has not been described previously. Since the NG2 glia depletion method described here is simple and versatile, it can be easily adapted to various physiological and neurological disease models, thereby enhancing our understanding of NG2 glia function both in the healthy and diseased brains.

PDGFRα mediated signaling is essential for the proliferation of NG2 glia in the embryonic brain and in primary NG2 glia cultures (Barres et al. 1992; Barres et al. 1993; Calver et al. 1998; Fruttiger et al. 1999). However, the role of PDGFRα for the postnatal maintenance of NG2 glia is unclear. Interestingly, a recent study found that PDGFRα also played important roles in the homeostatic control of oligodendroglia-lineage cells in the adult mouse brain (Dang et al. 2019). Here we show that inhibiting PDGF signaling promptly removes NG2 glia from long-term cultured brain slices, supporting a critical role for PDGF signaling on the maintenance of NG2 glia in the postnatal brain. These results raise possible safety issues of the clinical use of drugs targeting PDGFR signaling, many of which are currently being tested in clinical trials for various therapeutic purposes.

Microglia play vital roles in brain health and diseases. Depending on their cellular state, microglia exert distinct functions in different disease contexts (Sousa et al. 2018) and even in different stages of the same disease (Hamelin et al. 2018; Keren-Shaul et al. 2017). Understanding the mechanisms of the transition between different microglia cellular states is essential for developing therapeutic approaches against neurodegenerative diseases. Here we found that NG2 glia-microglia interaction played pivotal roles in regulating microglia states in the adult mouse brain. This finding may have profound implications for neurodegeneration. For example, it has been known that microglia associated with Aβ plaques in the AD brain display drastic downregulation of the homeostatic microglia signature genes (Keren-Shaul et al. 2017; Krasemann et al. 2017), which may be accompanied by functional changes that are directly related to disease progression. In contrast, Aβ plaque-associated NG2 glia showed a senescent phenotype (Zhang et al. 2019), indicating the loss of NG2 glia function in the plaque-containing microenvironment. The results of the current study suggest that the loss of the homeostatic signature in the plaque-associated microglia might be caused by a locally impaired NG2 glia-microglia interaction. This raises the possibility that targeting NG2 glia might be exploited for manipulating microglia states in neurodegenerative diseases.

We found that TGFβ signaling might be important in mediating the NG2 glia-microglia interaction, supporting a critical role for TGFβ in the regulation of microglia functions. TGFβ receptor 1 and 2, LTBP2, and many TGFβ-dependent microglia genes were profoundly downregulated upon depletion of NG2 glia. Previous studies have suggested that TGFβ pathway is important for normal microglia development and suppressed in microglia associated with neurodegenerative diseases (Butovsky et al. 2014; Krasemann et al. 2017). Disrupting TGFβ signaling in microglia through conditional removal of TGFβ receptor 2 in the adult brain resulted in loss of microglia homeostasis and altered microglia response to stimulations (Zoller et al. 2018). The action of TGFβ on microglia is highly selective and localized, with little spreading between neighboring cells (Qin et al. 2018). This process is non-cell-autonomous and relies on interactions between microglia and integrin αvβ8-bearing cells in the brain (Qin et al. 2018). High-throughput gene expression studies have suggested that NG2 glia express high levels of both subunits of integrin αvβ8 (Zhang et al. 2014), as well as TGFβs (Zhang et al. 2006). In this landscape of findings, the rapid and efficacious NG2 glia depletion method described here adds a useful component to the toolbox necessary to understand how NG2 glia regulate TGFβ signaling and maintains microglia in a homeostatic state.

## Materials and methods

### Ethics statement

All animal experiments were performed according to Swiss federal guidelines and were approved by the Animal Experimentation Committee of the Canton of Zurich (139/16 and ZH040-15). All efforts were made to minimize animal discomfort and suffering.

### Mouse experiments

C57BL/6J mice were purchased from Charles River, Germany. PDGFRα-CreER™ mice (Stock No: 018280) and inducible DTR (iDTR) mice (Stock No: 007900) were purchased from The Jackson Laboratory, USA. For tamoxifen induction, PDGFRα-CreER™, PDGFRα-CreER™/iDTR and iDTR mice were fed with tamoxifen-containing diet (ENVIGO RMS BV, TD.55125IC.I) for 4 weeks. For diphtheria toxin (DT) administration, the mice were injected with DT (Sigma, D0564) diluted in saline, and injected for five consecutive days (two injections per day, 200 ng per injection).

### Organotypic cultured slices

Cerebellar, hippocampal and cortical organotypic cultured slices were prepared according to a previously published protocol (Falsig and Aguzzi 2008). Briefly, brains from 12 days old (for cerebellar slices) or 7 days old (for hippocampal and cortical slices) C57BL/6J pups were cut into 350-μm thick sections in ice-cold Gey’s balanced salt solution (GBSS) supplemented with the glutamate receptor antagonist kynurenic acid (1 mM) and glucose with a vibratome (Leica). Slices with intact morphology were collected into the same GBSS solution and kept on ice. Six to eight slices were then put on a Millicell-CM Biopore PTFE membrane insert (Millipore) and kept on slice culture medium (50% vol/vol MEM, 25% vol/vol basal medium Eagle and 25% vol/vol horse serum supplemented with 0.65% glucose (w/vol), penicillin/streptomycin and glutamax (Invitrogen)) at 37 °C in cell culture incubator. The culture medium was changed every other day.

### PDGFR inhibitor treatment

PDGFR inhibitors CP673451 (HY-12050) and Crenolanib (HY-13223) were purchased from MedChemExpress, dissolved in dimethyl sulfoxide (DMSO, Sigma, 472301) and kept at −80 °C in aliquots. To achieve NG2 glia ablation, the inhibitors were supplemented in the slice culture medium. Fresh inhibitors were added every time when the culture medium was changed. The treatments started 14 days after the slice cultures were established with various concentrations and durations depending on the experiments. In all the PDGFR inhibitor treatment experiments, DMSO was used as control.

### Cell lines

The cell line Oli-neu (Jung et al. 1995) is a gift from Prof. Jacqueline Trotter in University of Mainz, Germany. Oli-neu cells were maintained in PLL-coated culture dishes in Opti-MEM medium supplemented with 1% inactivated horse serum, 1% Sato mix, 1% sodium pyruvate (ThermoFisher Scientific, 11360070), 1% GlutaMAX and penicillin/streptomycin. BV2 cell line was maintained in Opti-MEM medium supplemented with 5% FBS and penicillin/streptomycin. For CP673451 and Crenolanib treatment of Oli-neu and BV2 cells, the compounds were diluted in the culture medium at 500 nM concentration.

### Western blotting

To prepare samples for western blotting, COCS or brain tissues were lysed in RIPA buffer supplemented with proteinase inhibitor cocktail (Sigma, 000000011697498001). Western blotting was performed as previously described (Liu et al. 2015), with small modifications. As primary antibodies, we used mouse monoclonal antibody against actin (1:10000, Merck Millipore, MAB1501R), GFAP (1:2000, Sigma, AMAB91033), PLP (1:2000, Merck Millipore, MAB388); rabbit monoclonal antibody against NeuN (1:3000, Abcam, ab177487) and PDGFRβ (1:1000, Abcam, ab32570); rabbit polyclonal antibody against NG2 (1:1000, Millipore, AB5320), PDGFRα (1:500, Santa Cruz, SC-338) and Iba1 (1:1000, Wako, 019-19741). Appropriate HRP-conjugated secondary antibodies (1:10000, Jackson ImmunoResearch Laboratories) were used according to the host species of the primary antibodies. Membranes were visualized and digitized with ImageQuant (LAS-4000; Fujifilm). Optical densities of bands were analyzed by using ImageJ.

### Immunofluorescence

Immunofluorescence was performed according to procedures published previously (Falsig and Aguzzi 2008; Liu et al. 2018). Cultured brain slices were fixed in 4% PFA for 30 minutes and permeabilized with PBS supplemented with 0.1% Triton X-100 for 2 hours at room temperature; for mouse brain samples, mice were perfused with cold PBS and 4% PFA. After dissection, the brains were kept in 4% PFA for 4 hours, then transferred into 30% sucrose solution, and kept at 4 °C. After 3 day in sucrose solution, the brains were cut into 25-μm thick sections for staining. The following antibodies were used: anti-NG2 (1:500, a gift from Prof. Stallcup), anti-NG2 (1:500, Millipore, AB5320), anti-PDGFRα (1:500, a gift from Prof. Stallcup), anti-GFAP (1:300, Agilent Technologies, Z0334), anti-NeuN (1:500, Millipore, MAB377), anti-PDGFRβ (1:200, Abcam, ab32570), anti-CD13 (1:200, R&D Systems, AF2335-SP), anti-MBP (1:500, Millipore, NE1019), anti-CC1 (1:400, Millipore, OP80), anti-Iba1 (1:500, Wako, 019-19741), anti-CD68 (1:200, BioRad, MCA1957) and anti-TMEM119 (1:500, Synaptic Systems, 400 002). The cultured slices were incubated in the primary antibody for 3 days and brain sections were incubated in the primary antibody overnight at 4 °C. After washing, the cultured slices or brain sections were incubated in the secondary antibody overnight at 4 °C. Images were captured by using the FLUOVIEW FV10i confocal microscope (Olympus Life Science) and quantified with ImageJ.

### Quantitative real-time PCR (qRT-PCR)

qRT-PCR analysis was performed as previously described (Liu et al. 2018). Briefly, total RNA was extracted by using TRIzol (Invitrogen) according to the manufacturer’s instruction and cDNA was synthesized by using the QuantiTect Reverse Transcription kit (QIAGEN). qRT-PCR was performed using the SYBR Green PCR Master Mix (Roche) on a ViiA7 Real-Time PCR system (Applied Biosystems). The following primers were used: Mouse actin: sense 5′-AGATCAAGATCATTGCTCCTCCT-3′; antisense, 5′-ACGCAGCTCAGTAACAGTCC-3′. Mouse NG2: sense, 5′-ACCCAGGCTGAGGTAAATGC-3′; antisense, 5′-ACAGGCAGCATCGAAAGACA-3′. Mouse PDGFRα: sense, 5′-ATTAAGCCGGTCCCAACCTG-3′; antisense, 5′-AATGGGACCTGACTTGGTGC-3′. Mouse TNFα: sense, 5′-ACGTCGTAGCAAACCACCAA-3′; antisense, 5′-ATAGCAAATCGGCTGACGGT-3′. Mouse IL1β: sense, 5′-TGCAGCTGGAGAGTGTGGATCCC-3′; antisense, 5′-TGTGCTCTGCTTGTGAGGTGCTG-3′. Mouse IL6: sense, 5′-GCCTTCTTGGGACTGATGCT-3′; antisense, 5′-TGCCATTGCACAACTCTTTTCT-3′. Mouse IL12β: sense, 5′-TGGTTTGCCATCGTTTTGCTG-3′; antisense, 5′-ACAGGTGAGGTTCACTGTTTCT-3′. Mouse TGFβ1: sense, 5′-GTGGACCGCAACAACGCCATCT-3′; antisense, 5′-CAGCAATGGGGGTTCGGGCA-3′. Mouse TGFβ2: sense, 5′-CCGGAGGTGATTTCCATCTA-3′; antisense, 5′-GCGGACGATTCTGAAGTAGG-3′. Mouse CD11b: sense, 5′-AATTGGGGTGGGAAATGCCT-3′; antisense, 5′-TGGTACTTCCTGTCTGCGTG-3′. Mouse TMEM119: sense, 5′-CTTCACCCAGAGCTGGTTCCATA-3′; antisense, 5′-ATGATGAGGAAGGTCAGCGAG-3′. Mouse Olfml3: sense, 5′-CTAGCTGCCTTAGAGGAACGG-3′; antisense, 5′-CCTCCCTTTCAAGACGGTCC-3′. Mouse GPR34: sense, 5′-AGGAAAGCTTCAACTCAGTTCCT-3′; antisense, 5′-GAGCAAAGCCAGCTGTCAAC-3′. Mouse P2ry12: sense, 5′-TTGCACGGATTCCCTACACC-3′; antisense, 5′-GAAGCAGCCCCTTGGGTAAT-3′. Mouse Trem2: sense, 5′-CTGATCACAGCCCTGTCCCAAG-3′; antisense, 5′-TCTGACACTGGTAGAGGCCC-3′. Mouse Hexb: sense, 5′-CCAGACTGGAAGGTTGGTCC-3′; antisense, 5′-TGTAATATCGCCGAAACGCCT-3′. Mouse Tgfbr1: sense, 5′-ACGCGCTGACATCTATGCAA-3′; antisense, 5′-CCATCACTCTCAAGGCCTCA-3′. Mouse Tgfbr2: sense, 5′-CCGTGTGGAGGAAGAACGAC-3′; antisense, 5′-TGACAGCTATGGCAATCCCC-3′. Mouse Sall1: sense, 5′-AACTAAGCCGAGGACCAAGC-3′; antisense, 5′-CTCAAACATCAGCCGCTCAC-3′. Mouse Csf1r: sense, 5′-TCTTCCTCTGTTCCCTTTCAGG-3′; antisense, 5′-CCGATGCAGGTTGGAGAGTC-3′.

### RNA Sequencing

RNA sequencing of DMSO, CP673451 and Crenolanib treated slices was performed as previously described (Liu et al. 2018). Briefly, after 9 days of treatment, total RNA was extracted from the slices using the TRIzol™ Reagent (Invitrogen, 15596-018), further purified with RNeasy Plus universal mini kit (QIAGEN) and subjected to quality check using Bioanalyzer 2100 (Agilent Technologies) and Qubit (1.0) Fluorometer (Life Technologies). mRNA was enriched with Ribo-Zero Gold rRNA Removal Kit (Illumina) and libraries were prepared with the TruSeq RNA Sample Prep kit v2 (Illumina). Sequencing was performed using the TruSeq SBS kit v4-HS (Illumina) on Illumina Novaseq 6000 at 1 × 100 bp. Data analysis was performed by using the R and Bioconductor.

### Statistical analyses

Unless otherwise mentioned, unpaired, two-tailed student’s T-test was used for comparing data from two groups. All data were presented as mean ± SEM. Statistical analysis and data visualization were done by using GraphPad Prism 7 (GraphPad). P-values <0.05 were considered statistically significant.

## Acknowledgements

We thank Dr. William Stallcup for providing us with the polyclonal antibodies against NG2 and PDGFRα, Dr. Jacqueline Trotter for providing the cell line Oli-Neu, Dr. Annika Keller for providing us with the iDTR mice, the Functional Genomics Center Zurich for helping with RNA sequencing and basic data analyses, as well as Petra Schwarz, Julie Domange, Mirzet Delic and Olga Romashkina for excellent technical assistance. Dr. Asvin Lakkaraju, Dr. Assunta Senatore, Dr. Claudia Scheckel, Dr. Jiang-An Yin and Dr. Caihong Zhu read and commented the manuscript. A. Aguzzi is the recipient of an Advanced Grant of the European Research Council (ERC 250356) and is supported by grants from the Swiss National Foundation (SNF, including a Sinergia grant), the Swiss Initiative in Systems Biology, SystemsX.ch (PrionX, SynucleiX), the Klinische Forschungsschwerpunkte (KFSPs) “small RNAs” and “Human Hemato-Lymphatic Diseases”, and a Distinguished Investigator Award of the Nomis Foundation. The funders played no role in study design, data collection and analysis, decision to publish, or preparation of the manuscript.

## Legends for supplementary figures

**Suppl. fig 1.**
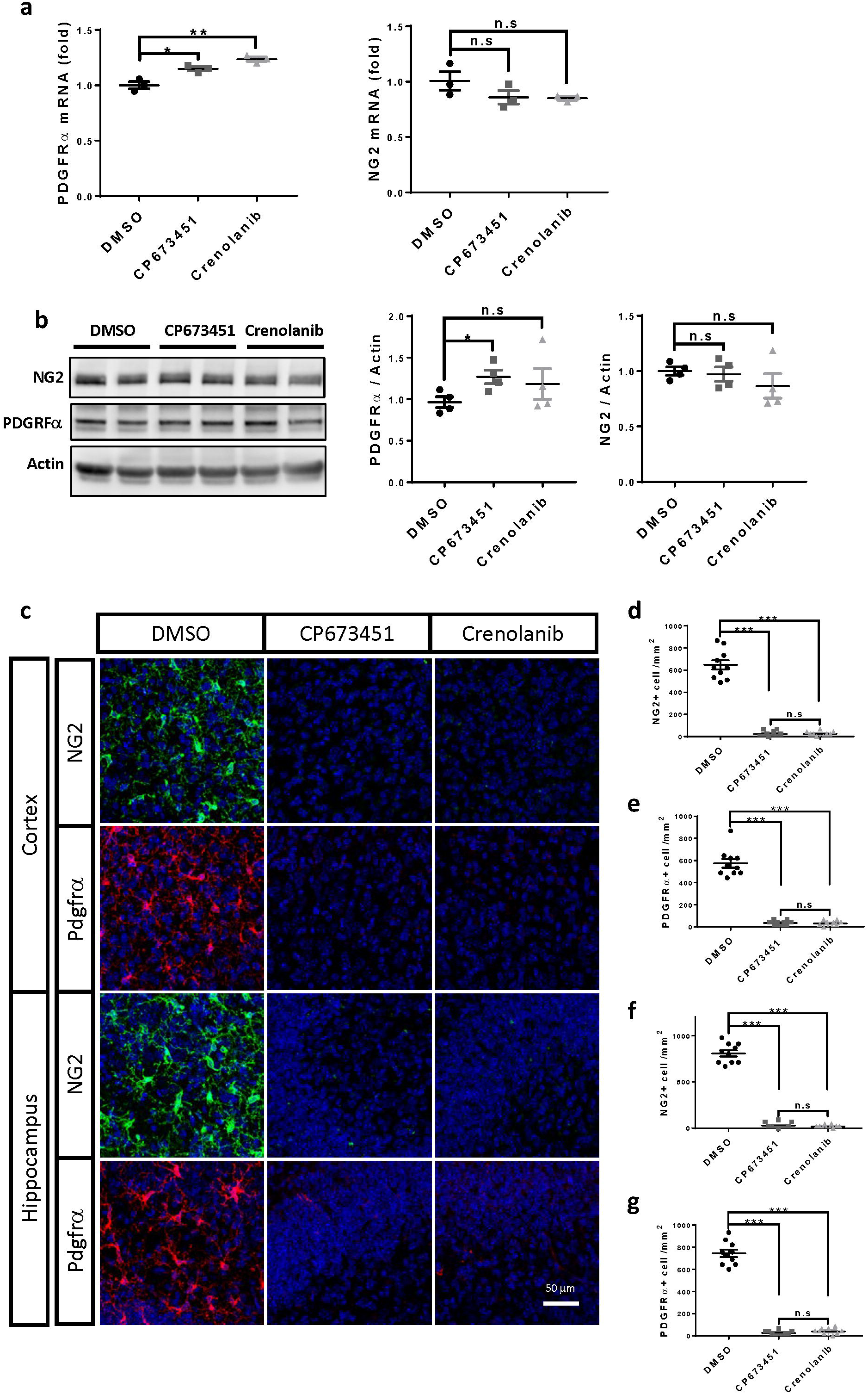
**a**, mRNA levels of NG2 and PDGFRα after CP673451 and Crenolanib treatment on Oli-neu cells. **b**, representative images of western blots and quantifications of NG2 and PDGFRα protein levels after CP673451 and Crenoilanib treatment on Oli-neu cells. **c**, immunofluorescence of PDGFRα and NG2 on hippocampal and cortical slices treated with 500 nM CP673451 and Crenolanib for 9 days. Scale bar: 50 μm. **d-g**, quantifications of PDGFRα^+^ and NG2^+^ cell numbers shown in **c**. * P<0.05; ** P<0.01; *** P< 0.001. n.s, not significant.

**Suppl. fig 2.**
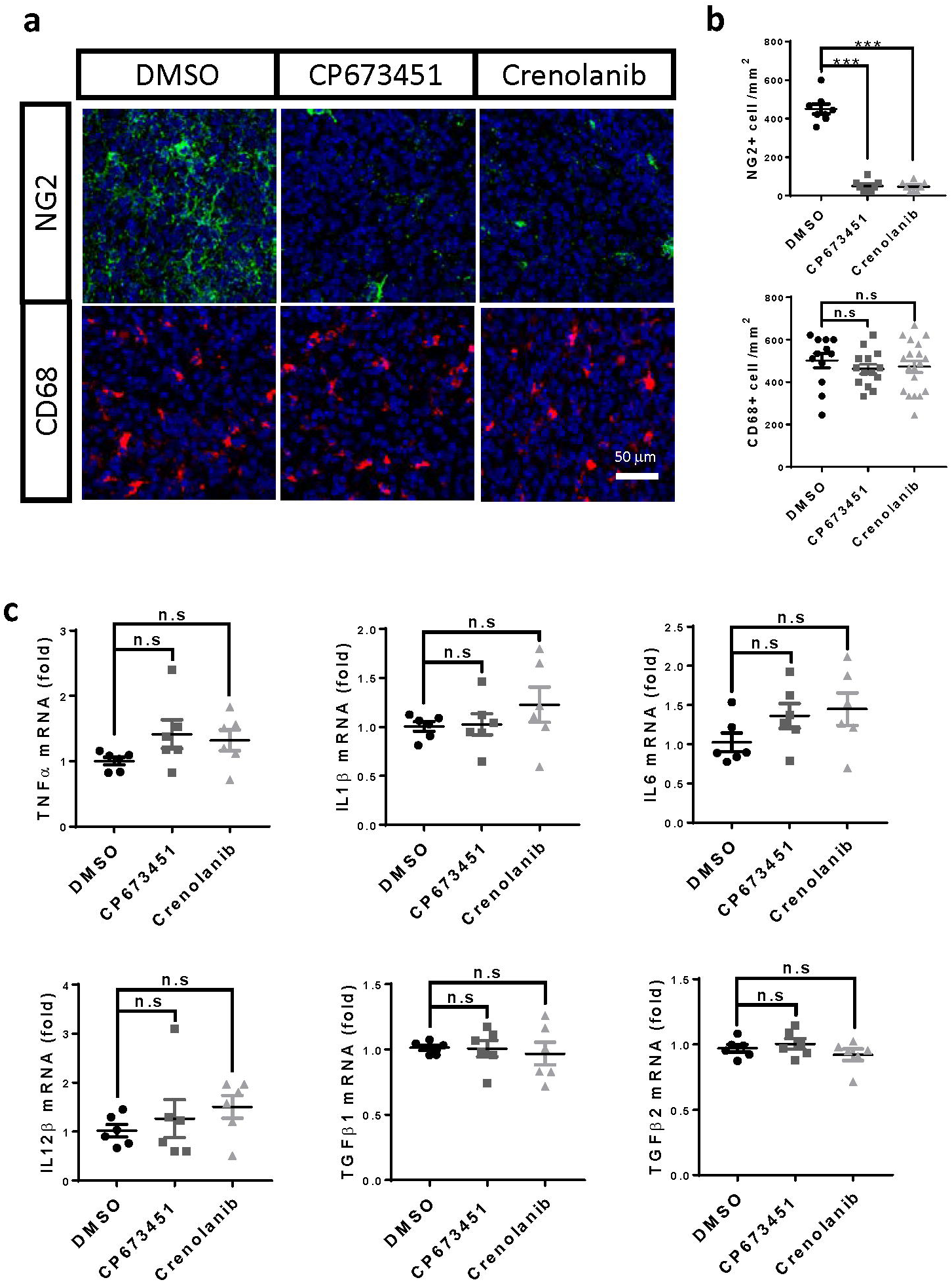
**a**, immunofluorescence of NG2 and CD68 on slices treated with 500 nM CP673451 and Crenolanib for 9 days. Scale bar: 50 μm. **b**, quantifications of NG2^+^ or CD68^+^ cell numbers shown in **a. c**, mRNA levels of TNFa, IL1β, IL6, IL12β, TGFβ1 and TGFβ2 after NG2 glia depletion. *** P< 0.001. n.s, not significant.

**Suppl. fig 3.**
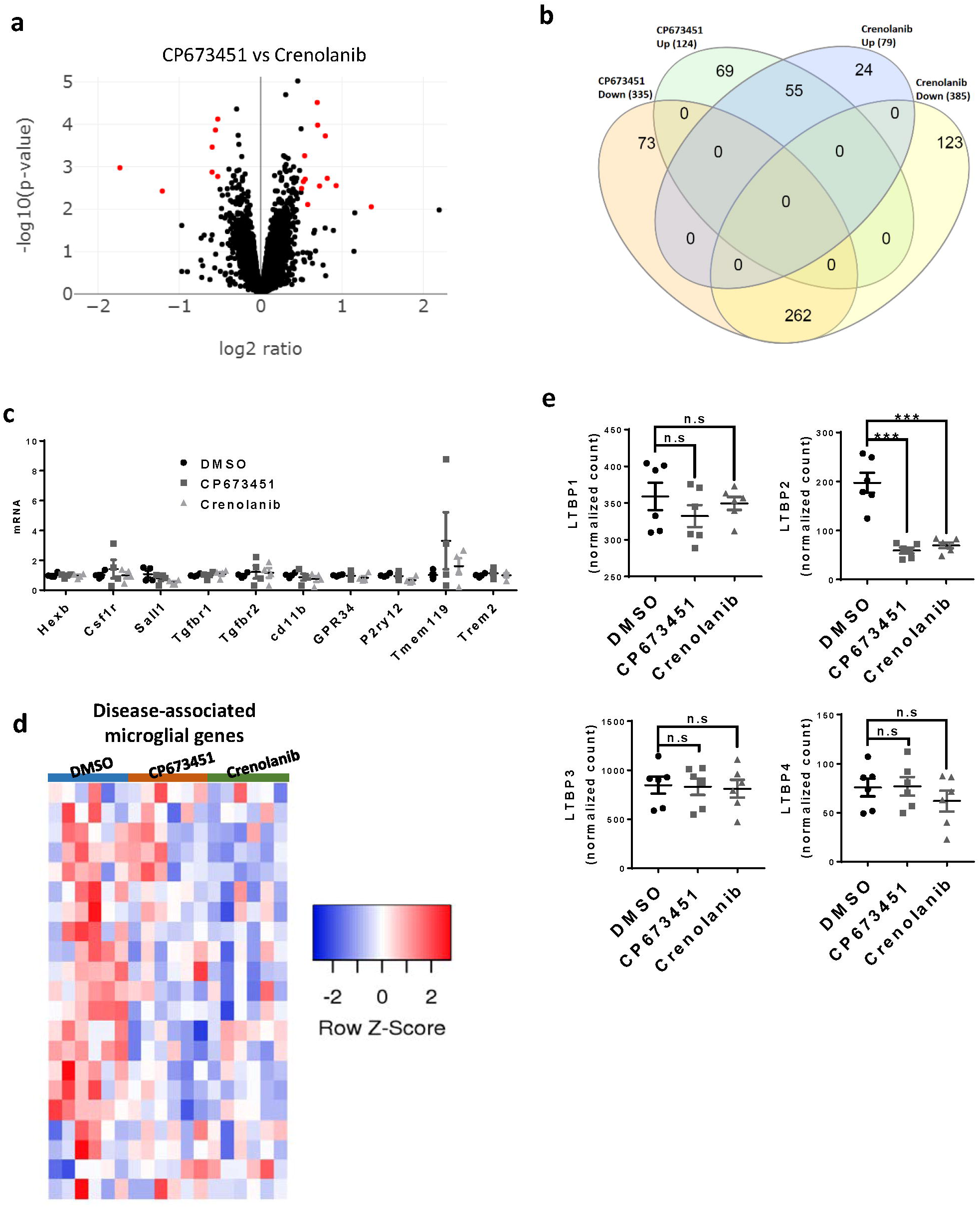
**a**, volcano plot of RNAseq data comparing CP673451 and Crenolanib treated brain slices. **b**, Venn diagram showing the overlapping genes between different experiment groups. **c**, mRNA levels of a subset of homeostatic microglia signature genes after CP673451 and Crenolanib treatment on BV2 microglia. **d**, heatmap of disease-associated microglia genes after NG2 glia depletion. **e**, expression levels of LTBPs in the RNAseq dataset.

**Suppl. fig 4.**
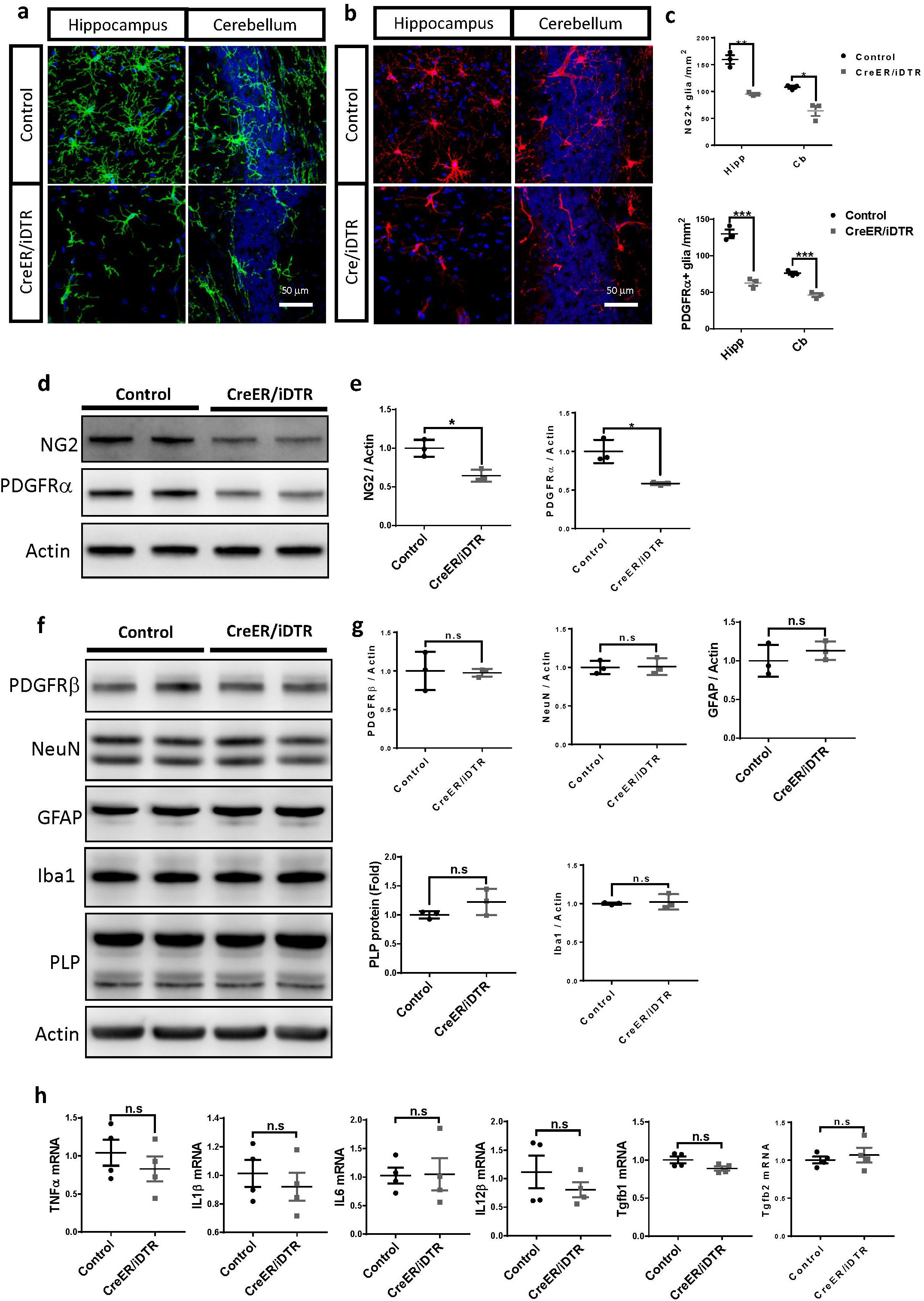
**a**, immunofluorescence of NG2 in hippocampus and cerebellum of PDGFR-CreER™/iDTR mice and control mice. Scale bar: 50 μm. **b**, immunofluorescence of PDGFRα in hippocampus and cerebellum of PDGFR-CreER™/iDTR mice and control mice. Scale bar: 50 μm. **c**, quantifications of NG2^+^ and PDGFRα^+^ cell numbers shown in **a** and **b. d**, representative images of western blots of NG2 and PDGFRα in the brains of PDGFR-CreER™/iDTR and control mice. **e**, quantifications of NG2 and PDGFRα protein levels shown in **d. f**, representative images of western blots of PDGFRβ, NeuN, GFAP, Iba1 and PLP in the brains of PDGFR-CreER™/iDTR and control mice. **g**, quantifications of protein levels shown in **f. h**, mRNA levels of inflammatory genes in the brains of PDGFR-CreER™/iDTR and control mice after DT injection. * P<0.05; ** P<0.01; *** P< 0.001. n.s, not significant.

